## Supplementary figures and images for "Genome-wide screen for deficiencies modifying *Cyclin G*-induced developmental instability in *Drosophila melanogaster*"

### Supplementary Figure 1

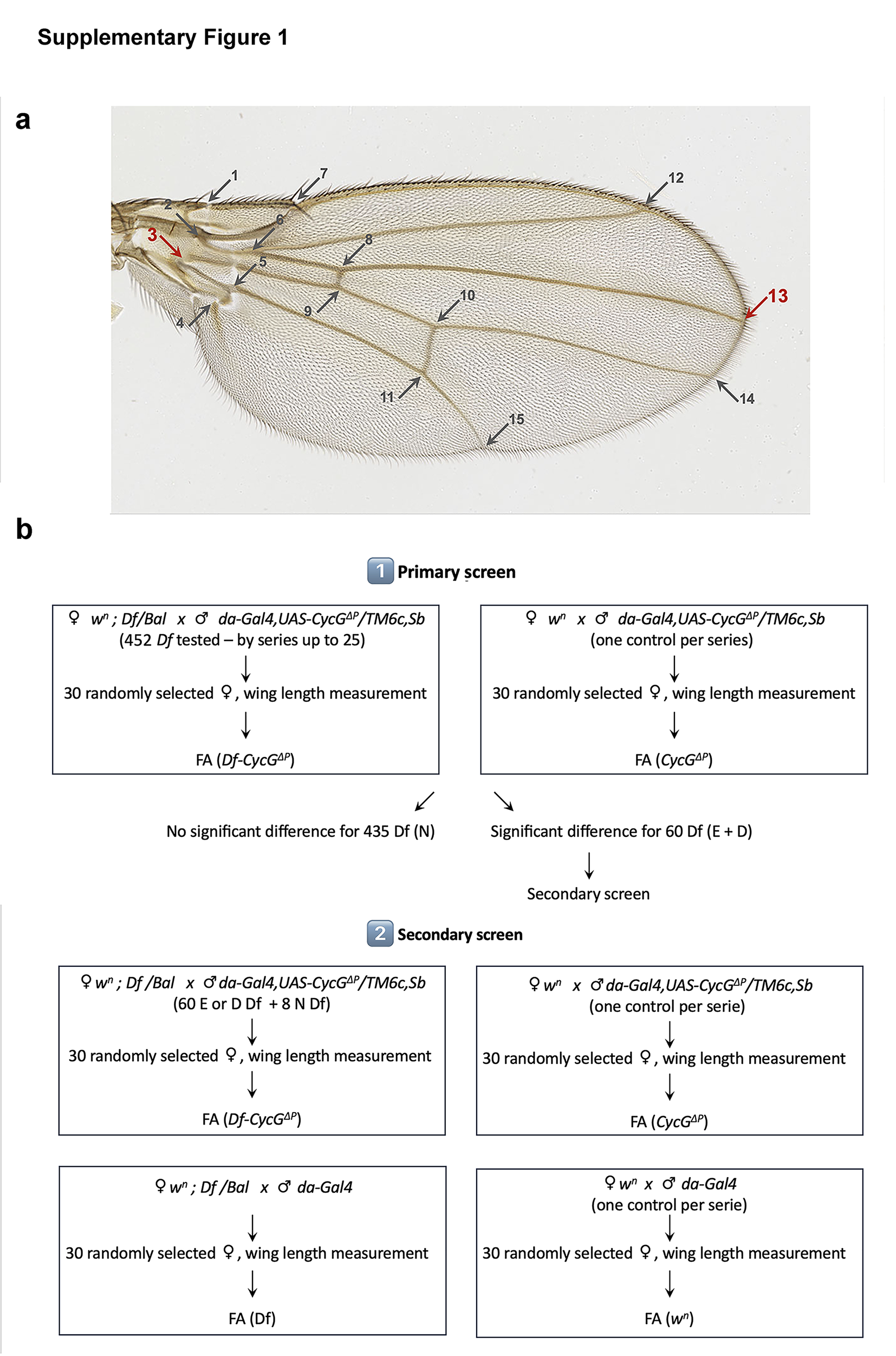

### Supplementary Figure 2

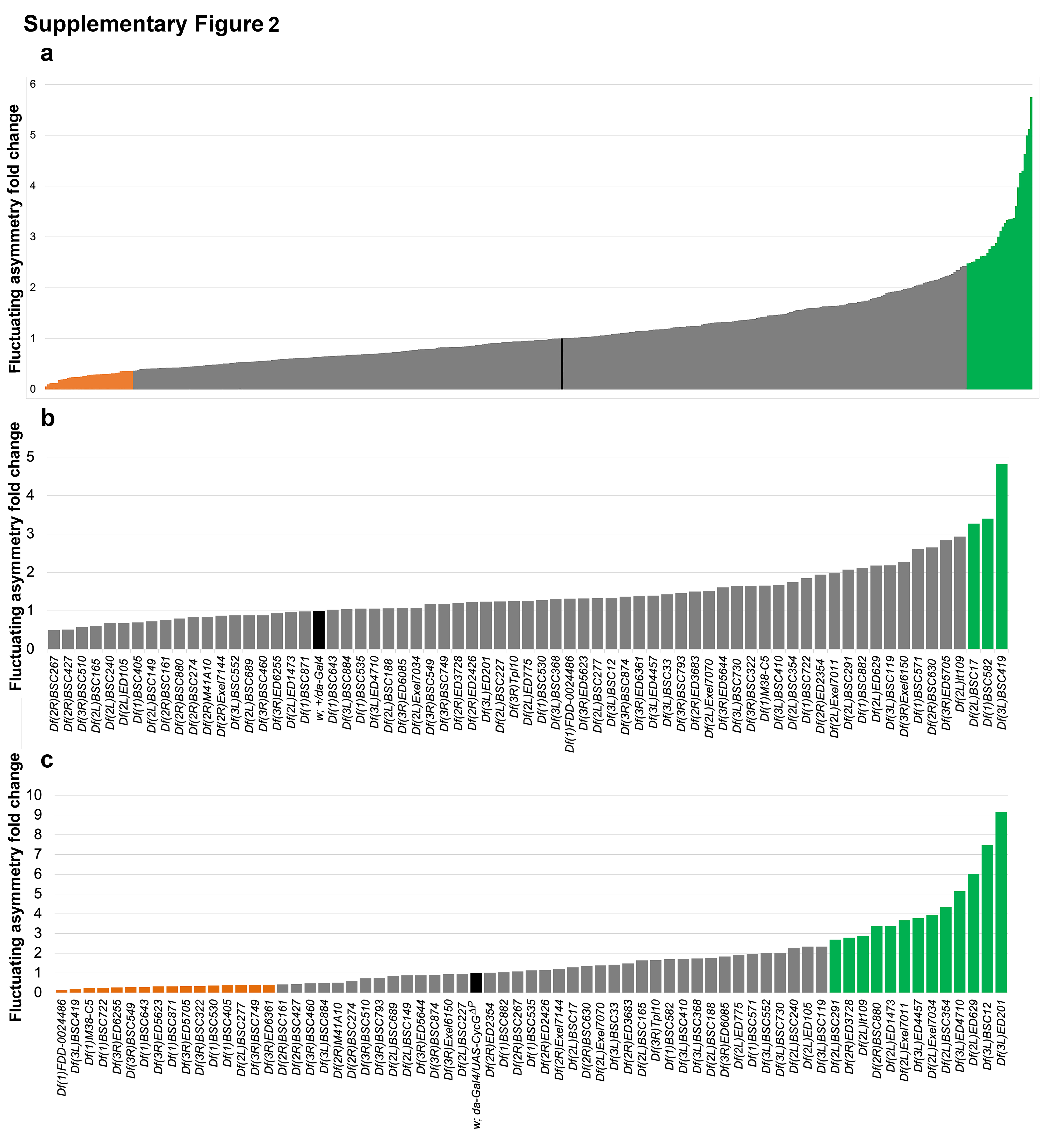

### Supplementary Figure 3

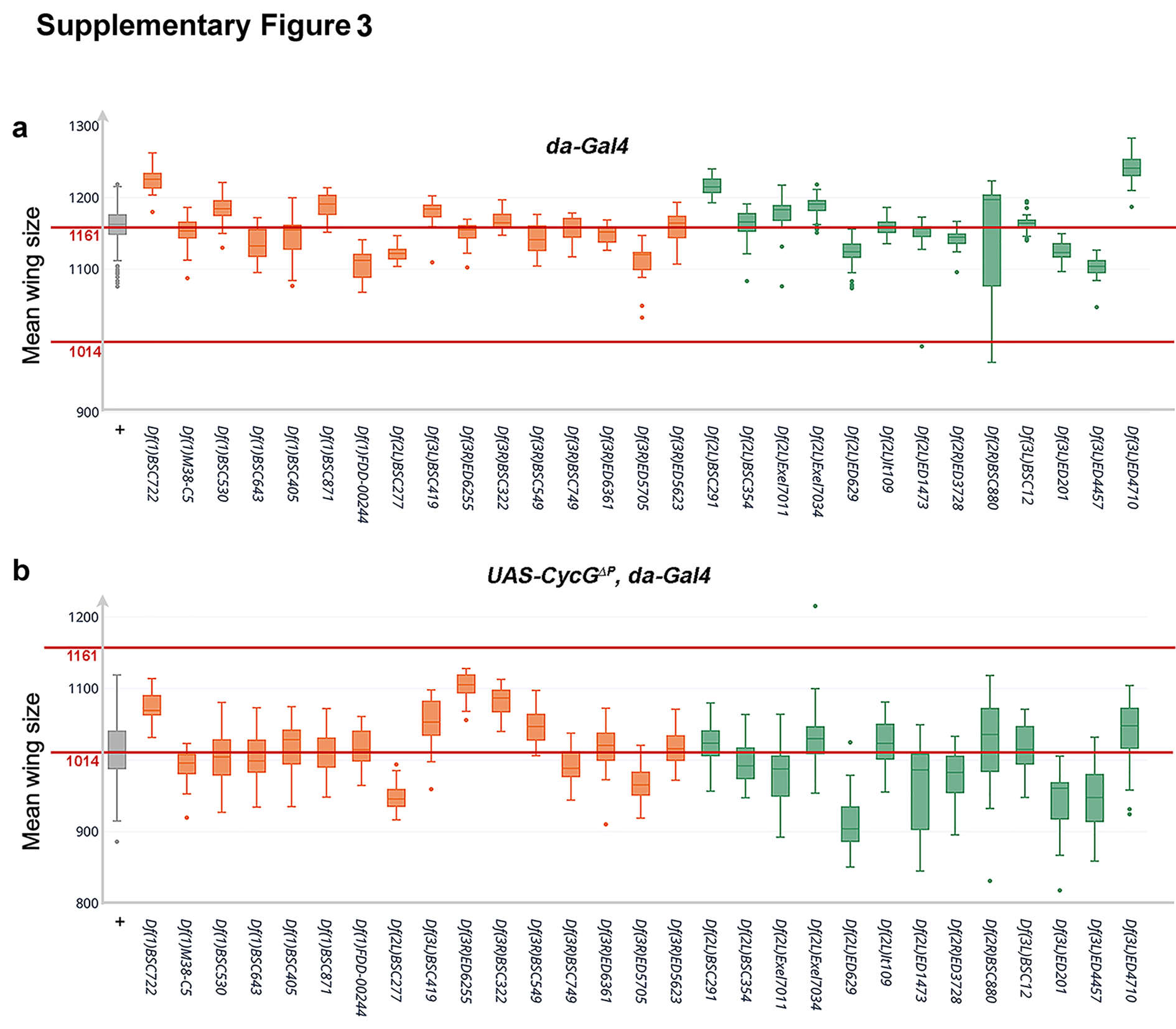
